# Exceptional subgenome stability and functional divergence in allotetraploid teff, the primary cereal crop in Ethiopia

**DOI:** 10.1101/580720

**Authors:** Robert VanBuren, Ching Man Wai, Jeremy Pardo, Alan E. Yocca, Xuewen Wang, Hao Wang, Srinivasa R. Chaluvadi, Doug Bryant, Patrick P. Edger, Jeffrey L. Bennetzen, Todd C. Mockler, Todd P. Michael

## Abstract

Teff (*Eragrostis tef*) is a cornerstone of food security in the Horn of Africa, where it is prized for stress resilience, grain nutrition, and market value. Despite its overall importance to small-scale farmers and communities in Africa, teff suffers from low production compared to other cereals because of limited intensive selection and molecular breeding. Here we report a chromosome-scale genome assembly of allotetraploid teff (variety ‘Dabbi’) and patterns of subgenome dynamics. The teff genome contains two complete sets of homoeologous chromosomes, with most genes maintained as syntenic gene pairs. Through analyzing the history of transposable element activity, we estimate the teff polyploidy event occurred ∼1.1 million years ago (mya) and the two subgenomes diverged ∼5.0 mya. Despite this divergence, we detected no large-scale structural rearrangements, homoeologous exchanges, or bias gene loss, contrasting most other allopolyploid plant systems. The exceptional subgenome stability observed in teff may enable the ubiquitous and recurrent polyploidy within Chloridoideae, possibly contributing to the increased resilience and diversification of these grasses. The two teff subgenomes have partitioned their ancestral functions based on divergent expression patterns among homoeologous gene pairs across a diverse expression atlas. The most striking differences in homoeolog expression bias are observed during seed development and under abiotic stress, and thus may be related to agronomic traits. Together these genomic resources will be useful for accelerating breeding efforts of this underutilized grain crop and for acquiring fundamental insights into polyploid genome evolution.

## Introduction

Thirty crop species supply over 90% of the world’s food needs and this narrow diversity reduces global food security. Humans have domesticated several hundred distinct plant species, but most are underutilized, under-improved, and restricted to their regions of origin ^1^. Although food systems have become increasingly diverse in the last few decades, many locally adapted species have been replaced by calorically dense staple crops, resulting in global homogeneity ^2^. Many underutilized and “orphan” crop species have desirable nutritional profiles, abiotic and biotic stress resilience, and untapped genetic potential for feeding the growing population under the changing climate.

Teff is the staple grain crop in Ethiopia, and it is preferred over other cereals because of its nutritional profile, low input demand, adaptability, and cultural significance. Unlike other major cereals, teff is grown primarily by small-scale, subsistence farmers. An estimated 130,000 locally adapted cultivars have been developed. Teff is among the most resilient cereals, tolerating marginal and semi-arid soils that are unsuitable for wheat, maize, sorghum, and rice production. Teff was likely domesticated in the northern Ethiopian Highlands where much of the genetic diversity can be found ^3-5^. Consistent yields of small, nutritious seeds were the primary domestication targets of teff, contrasting most cereals where large seed heads and high productivity under tillage were desirable ^5^. Despite its stress tolerance, yield improvements lag behind other cereals because of issues related to lodging, seed shattering, extreme drought, and poor agronomic practices ^6^. Teff and other orphan cereals have undergone limited intensive selection for high productivity under ideal conditions, and rapid gains should be possible with advanced breeding and genome selection. A draft genome is available for the teff cultivar ‘Tsedey’ (DZ-Cr-37) ^7^, but the utility of this reference is limited given its fragmented and incomplete nature.

The wild progenitor of teff is likely *Eragrostis pilosa*; a hardy wild grass sharing considerable overlap in morphological, genetic, and karyotype traits with teff ^8,9^. *E. tef* and *E.pilosa* are allotretraploids that arose from a shared polyploidy event of merging two distant, unknown diploid genomes ^9^. Many crop plants are polyploid, and genome doubling can give rise to emergent traits such as spinnable fibers in cotton ^10^, morphological diversity in Brassica sp. ^11^, and new aromatic profiles of strawberry fruits ^12^. Successful establishment of allopolyploids requires coordination of two distinct sets of homoeologous genes and networks, and often a ‘dominant’ subgenome emerges to resolve genetic and epigenetic conflicts ^13-15^. The effect of polyploidy on desirable traits and interactions between the two subgenomes remains untested in teff. Polyploidy is found in more than 90% of species within the grass subfamily containing teff (Chloridoideae), and this has been hypothesized to contribute to the stress tolerance and diversification of these grasses ^16^. Here, we report a chromosome-scale assembly of the teff A and B subgenomes and test for patterns of subgenome interactions and divergence.

## Results

### Genome assembly and annotation

We built a chromosome-scale assembly of the allotetraploid teff genome using a combination of long read SMRT sequencing and long-range high-throughput chromatin capture (Hi-C). In total, we generated 5.5 million filtered PacBio reads collectively spanning 52.9 Gb or 85x coverage of the estimated 622 Mb ‘Dabbi’ teff genome. PacBio reads were error corrected and assembled using Canu^17^ and the resulting contigs were polished to remove residual errors with Pilon^18^ using high coverage Illumina data (45x). The PacBio assembly has a contig N50 of 1.55 Mb across 1,344 contigs with a total assembly size of 576 Mb; 92.6% of the estimated genome size. The graph-based structure of the assembly has few bubbles corresponding to heterozygous regions between haplotypes but contains numerous ambiguities related to high copy number long terminal repeat (LTR) retrotransposons (Supplemental Figure 1). This pattern was also observed in the genome assembly graph a the closely related grass, *Oropetium thomaeum* ^19^. The average nucleotide identity between homoeologous regions in teff is 93.9% in protein coding regions. Thus, high sequence divergence facilitated accurate phasing and assembly. We utilized twenty random fosmids to assess the accuracy of the PacBio-based assembly (Supplemental Table 1). The fosmids collectively span 351kb and have an average identity of 99.9% to the teff genome with individual fosmids ranging from 99.3 to 100%. This suggests that our assembly is mostly complete and accurately polished.

Contigs from the Canu based draft genome were anchored into a chromosome-scale assembly using a Hi-C based scaffolding approach. Illumina reads from the Hi-C library were aligned to the PacBio contigs with BWA ^20^ followed by proximity based clustering using the Juicer pipeline ^21^. 150bp paired-end reads and aggressive filtering of non-uniquely mapped reads were used to minimize chimeric mapping errors between homoeologous regions. After filtering, twenty high-confidence clusters were identified, consistent with the haploid chromosome number of teff (2n=40; Figure 1). In total, 687 contigs collectively spanning 96% of the assembly (555 Mb) were anchored and oriented across the 20 pseudomolecules (Table 1). Pseudomolecules ranged in size from 19 to 40 Mb, consistent with the teff karyotype ^22^. Seven chimeric contigs corresponding to joined telomeres were identified and split based on Hi-C interactions. As described in the accompanying manuscript (see Wang et al. 2019), this genome assembly was compared to a detailed genetic map of teff to revise and confirm chromosome-scale assemblies for all 20 teff chromosomes, thus providing the opportunity to discover the A and B genomes from the diploid progenitors of this allotetraploid (see below).

**Table 1.**
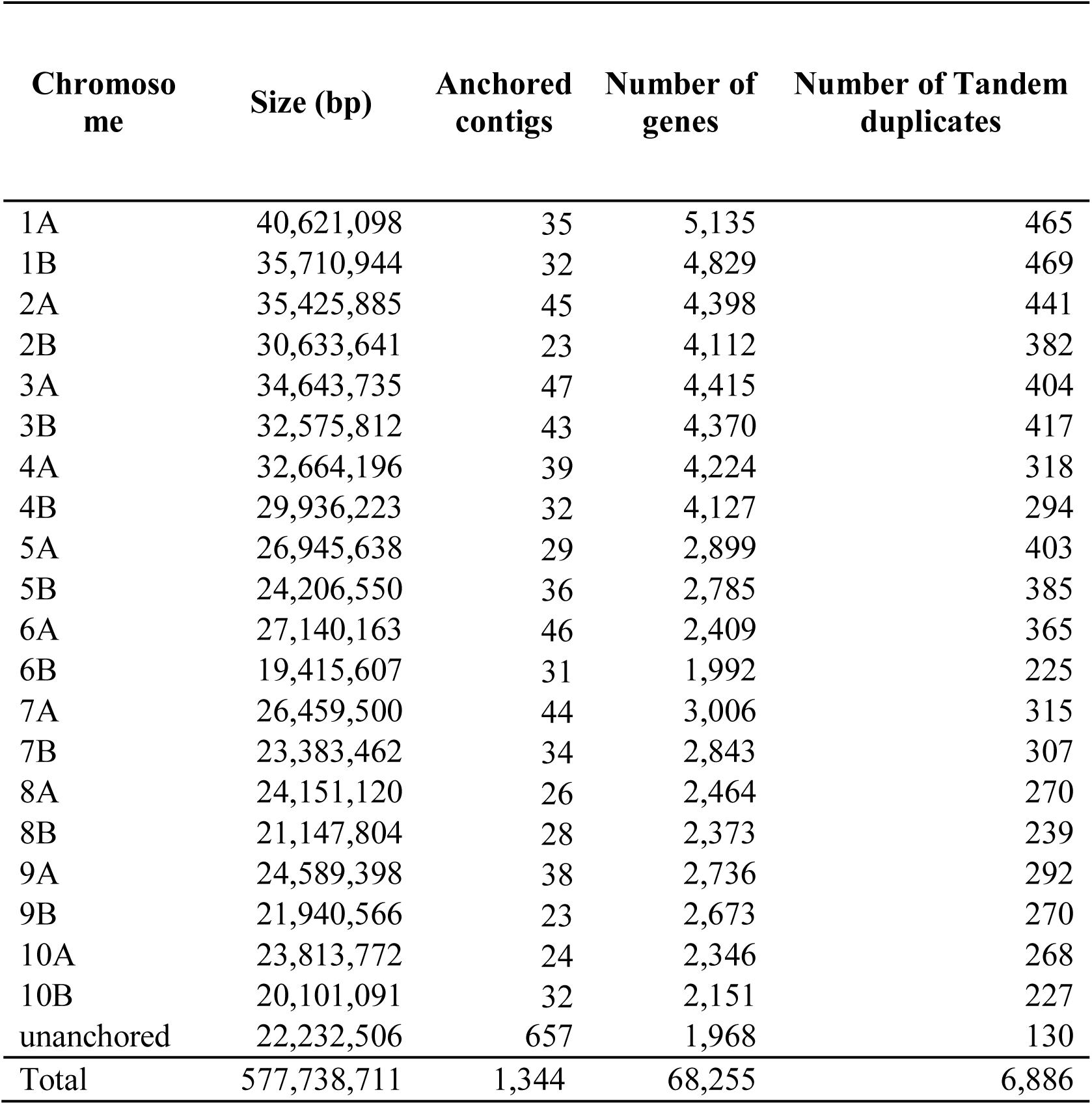
Summary statistics of the teff genome

**Figure 1.**
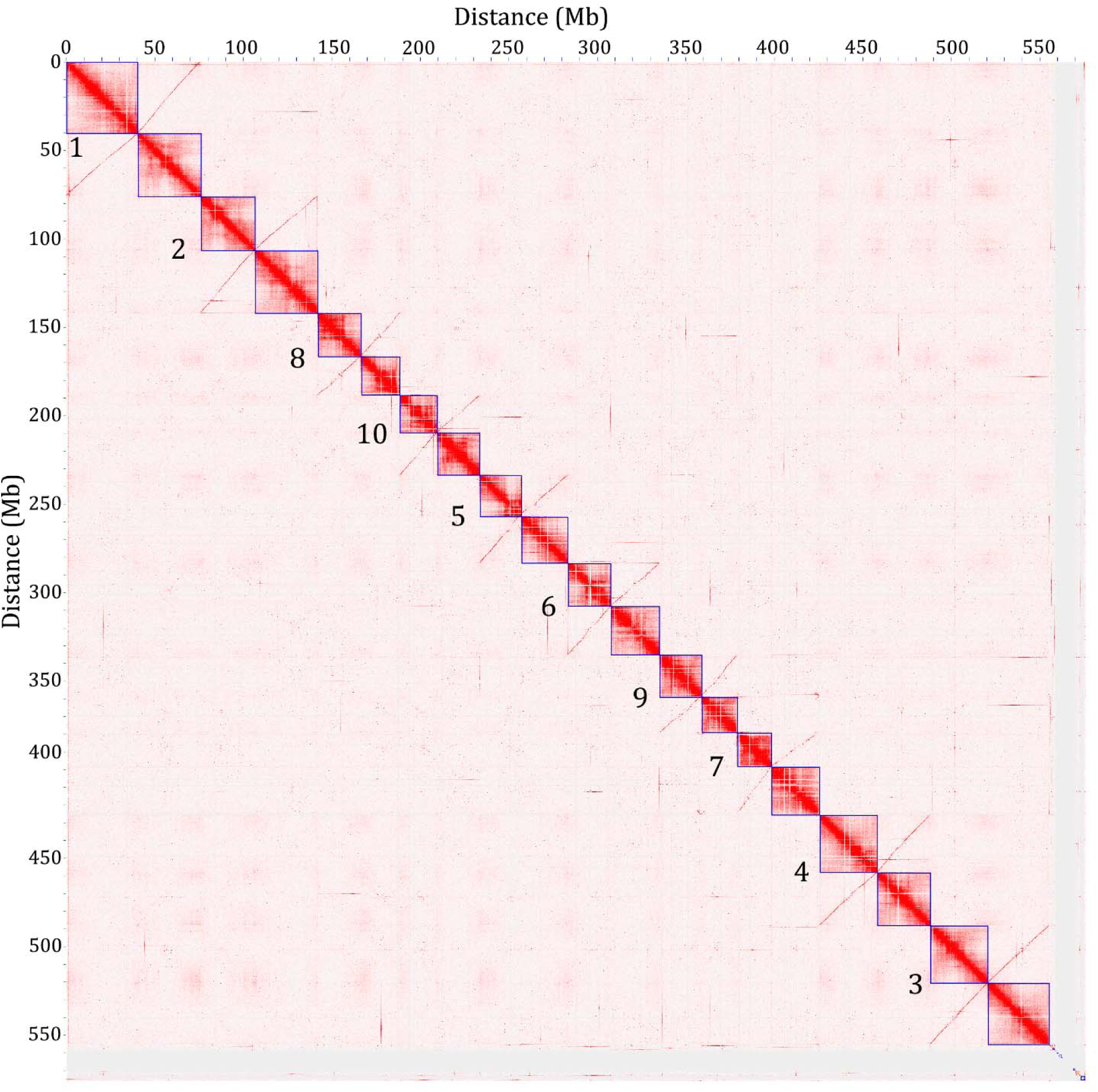
Hi-C based clustering of the teff genome. Heat map showing the density of Hi-C interactions between contigs with red indicating high density of interactions. Distinct chromosomes are highlighted by blue boxes and homoeologous chromosome pairs are numbered.

The teff genome was annotated using the MAKER pipeline. Transcript support from a large-scale expression atlas and protein homology to Arabidopsis and other grass genomes were used as evidence for *ab initio* gene prediction. After filtering transposon-derived sequences, *ab initio* gene prediction identified 68,255 gene models. We assessed the annotation quality using the Benchmarking Universal Single-Copy Ortholog (BUSCO) Embryophyta dataset. The annotation contains 98.1% of the 1,440 core Embryophyta genes and the majority (1,210) are found in duplicate in the A and B subgenomes.

The teff cultivar ‘Tsedey’ (DZ-Cr-37) was previously sequenced using an Illumina based approach, yielding a highly fragmented draft genome with 14,057 scaffolds and 50,006 gene models ^7^. The fragmented nature of this assembly and incomplete annotation hinders downstream functional genomics, genetics, and marker-assisted breeding of teff. We compared the ‘Tsedey’ assembly with our ‘Dabbi’ reference to identify cultivar-specific genes and differences in assembly quality. Only 30,424 (60.8%) of the ‘Tsedey’ gene models had homology (>95% sequence identity) to gene models in our ‘Dabbi’ reference, including 9,866 homoeologous gene pairs. Only 20,208 (29.6%) of our ‘Dabbi’ gene models had homology to ‘Tsedey’ gene models. The remaining gene models were unannotated or unassembled in the ‘Tsedey’ assembly. Only one-third of the ‘Tsedey’ genome is assembled into scaffolds large enough to be classified as syntenic blocks to ‘Dabbi’, which is an unavoidable artifact of the poor assembly quality and low contiguity. Because of the fragmented nature of the ‘Tsedey’ assembly, we were unable to identify lineage-specific genes. Hence, the genomic resources presented here represent a significant advance over previous efforts.

### Origins and subgenome dynamics

Teff is an allotetraploid with unknown diploid progenitors, but the polyploidy event is likely shared with other closely related *Eragrostis* species ^9^. Because the diploid progenitors are unknown and possibly extinct, we utilized the centromeric array sequences to distinguish the homoeologous chromosomes from the A and B subgenomes of teff. Centromeric (CenT) repeat arrays in teff range from 3.7 kb to 326 kb in size for each chromosome and individual arrays contain 22 to 824 copies (Supplemental Table 2). We identified two distinct CenT arrays in teff (hereon referend to as CenTA and CenTB). CenTA and CenTB are the same length (159 bp) but have different sequence composition (Supplemental Figure 2b). Alignment of the consensus CenT arrays identified several distinguishing polymorphisms and a maximum likelihood phylogenetic tree separated the CenT arrays into two well-supported clades (Supplemental Figure 2a). Each clade contains one member from each of the ten homoeologous chromosome pairs and this classification likely represents differences in Cen array composition between the diploid progenitor species. This approach allowed us to accurately distinguish homoeologous chromosome pairs from the A and B subgenomes and verifies the allopolyploid origin of teff.

The Teff subgenomes have 93.9% sequence homology in the coding regions, suggesting that either the polyploidy event was relatively ancient or that the progenitor diploid species were highly divergent ^23^. To estimate the divergence time of the A and B subgenomes, we calculated Ks (synonymous substitutions per synonymous site) between homoeologous gene pairs. Teff homoeologs have a single Ks peak with a median of 0.15 (Supplemental Figure 3), corresponding to a divergence time of ∼5 million years based on a widely used mutation rate for grasses ^24^. The ten pairs of homoeologous chromosomes are highly syntenic with no large-scale structural rearrangements. The A subgenome is 13% (37 Mb) larger in size but contains only 5% more genes than the B subgenome (34,032 vs. 32,255; Table 1). Most genes (54,846) are maintained as homoeologous pairs and 13,409 are found in only one subgenome. We identified 6,876 tandemly duplicated genes with array sizes ranging from 2 to 15 copies. Of the 2,748 tandem arrays, 998 are found in both subgenomes, while 864 and 1,008 occur in only the A and B subgenomes, respectively (Table 1). Copy number varies extensively in shared arrays between the subgenomes.

The monoploid genome size of teff is relatively small (∼300 Mb) compared to other polyploid grasses, and repetitive elements constitute a low percentage (25.6%) of the genome. Long terminal repeat retrotransposons (LTR-RTs) are the most abundant repetitive elements, spanning at least 115.9 Mb or ∼20.0% of the genome (Supplemental Table 3). This predicted percentage is somewhat lower than that reported for other small grass genomes such as Oropetium (250 Mb; 27%) ^19,25^ and Brachypodium (272 Mb; 21.4%) ^26^. We classified LTRs into families and compared their abundance and insertion times (Figure 2). A particular window of activity was seen for six families of LTR-RTs that were active only in the A genome progenitor or the B genome progenitor (Supplemental Figure 4, Supplemental Table 4). The insertion times for these genome-specific LTR-RTs were all greater than 1.1 mya, indicating the two subgenomes were evolving independently during this period. Hence, this LTR-RT analysis both confirms the A and B genome designations, and provides a novel methodology for determining the date of polyploid formation. In teff, these data indicate that the ancestral polyploidy was established ∼1.1 mya.

**Figure 2.**
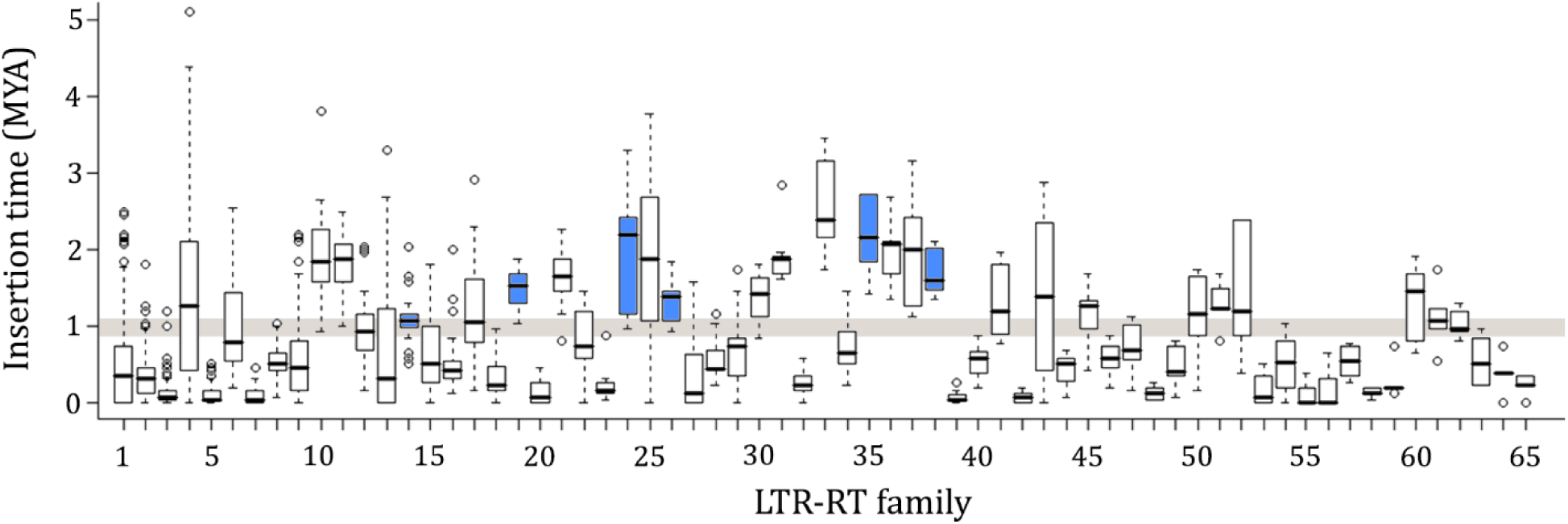
Insertion dynamics of 65 LTR-RT families in teff. Box plots of insertion time for the 65 LTR-RT families having ≥ 5 intact LTR elements are plotted. Families 1-5 have ≥ 100 intact LTRs, 6-33 have ≥ 10 LTRs, and 34-65 have ≥ 5 LTRs. The six subgenome specific families are highlighted in blue and the estimated range for the teff polyploidy event is shown in brown. A substitution rate of 1.3e-8 per site per year was used to infer the element insertion times.

Five of the six subgenome-specific LTR-RT families were found only in the A subgenome, suggesting that LTR-RTs accumulate more rapidly in the A subgenome or are purged more effectively in the B subgenome. This recent bursts of LTR-RT activity contributes to the 13% larger size of the A subgenome. There are 24 families with median insertion times between 1.1 and 2.4 MYA, and the remaining 18 families do not exhibit subgenomic specificity. Of these, 15 show no apparent burst in amplification, and three evidence of very recent (post-polyploid) activity (Figure 4, Supplementary Figure 4, Supplemental Table 5).

**Figure 3.**
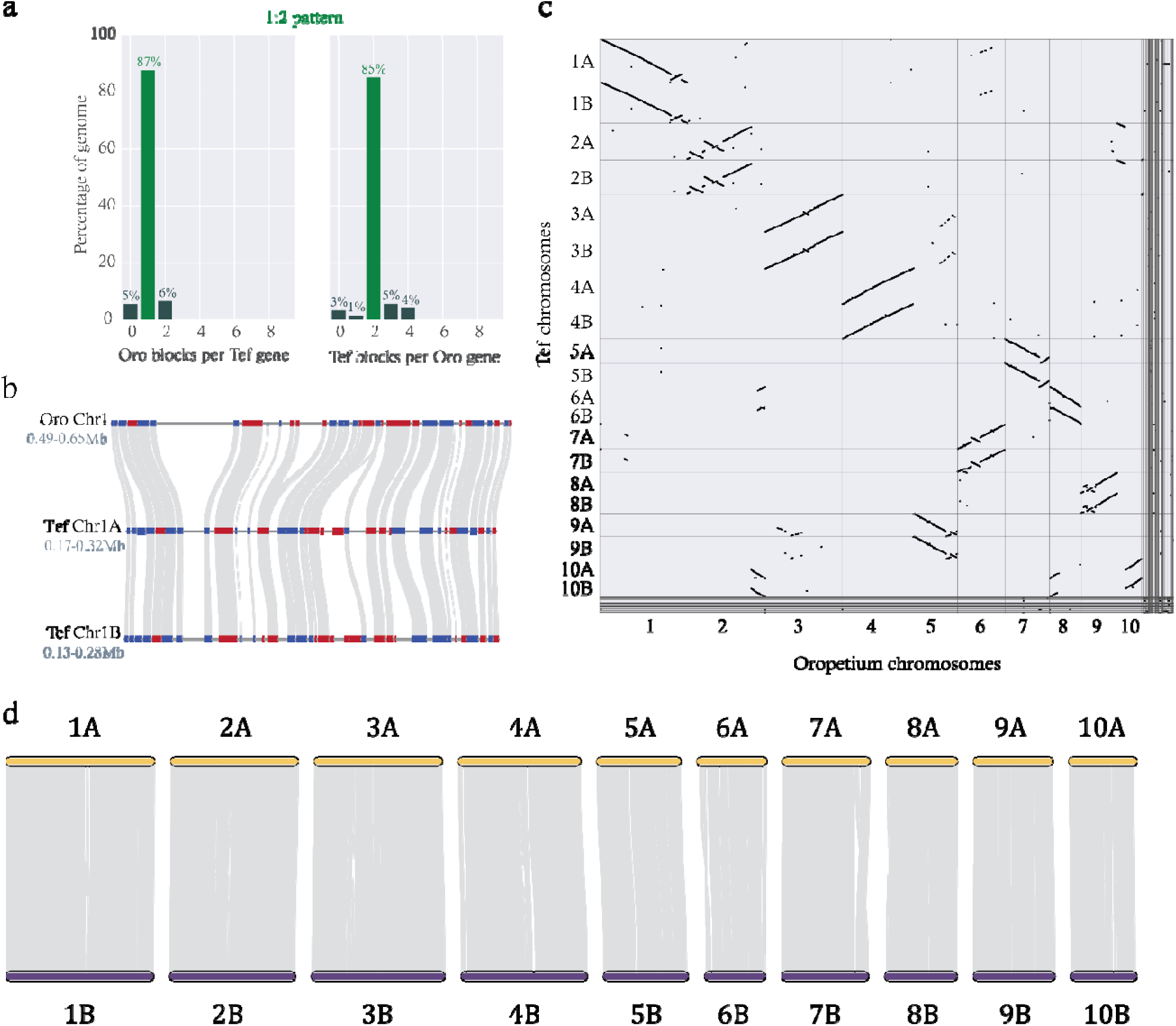
Comparative genomics of the teff genome. (a) Ratio of syntenic depth between Oropetium and teff. Syntenic blocks of Oropetium per teff gene (left) and syntenic blocks of teff per Oropetium gene (right) are shown indicating a clear 1:2 pattern of Oropetium to teff. (b) Microsynteny of the teff and Oropetium genomes. A region of the Oropetium Chromosome 1 and the corresponding syntenic regions in homoeologous teff Chromosomes 1 A and B are shown. Genes are shown in red and blue (for forward and reverse orientation respectively) and syntenic gene pairs are connected by grey lines. (c) Macrosynteny of the teff and Oropetium genomes. Syntenic gene pairs are denoted by gray points. (d) Collineariy of the teff subgenomes. The ten chromosomes belonging to the teff A and B subgenomes are shown in yellow and purple respectively. Syntenic blocks between homoeologous regions are shown in grey.

**Figure 4.**
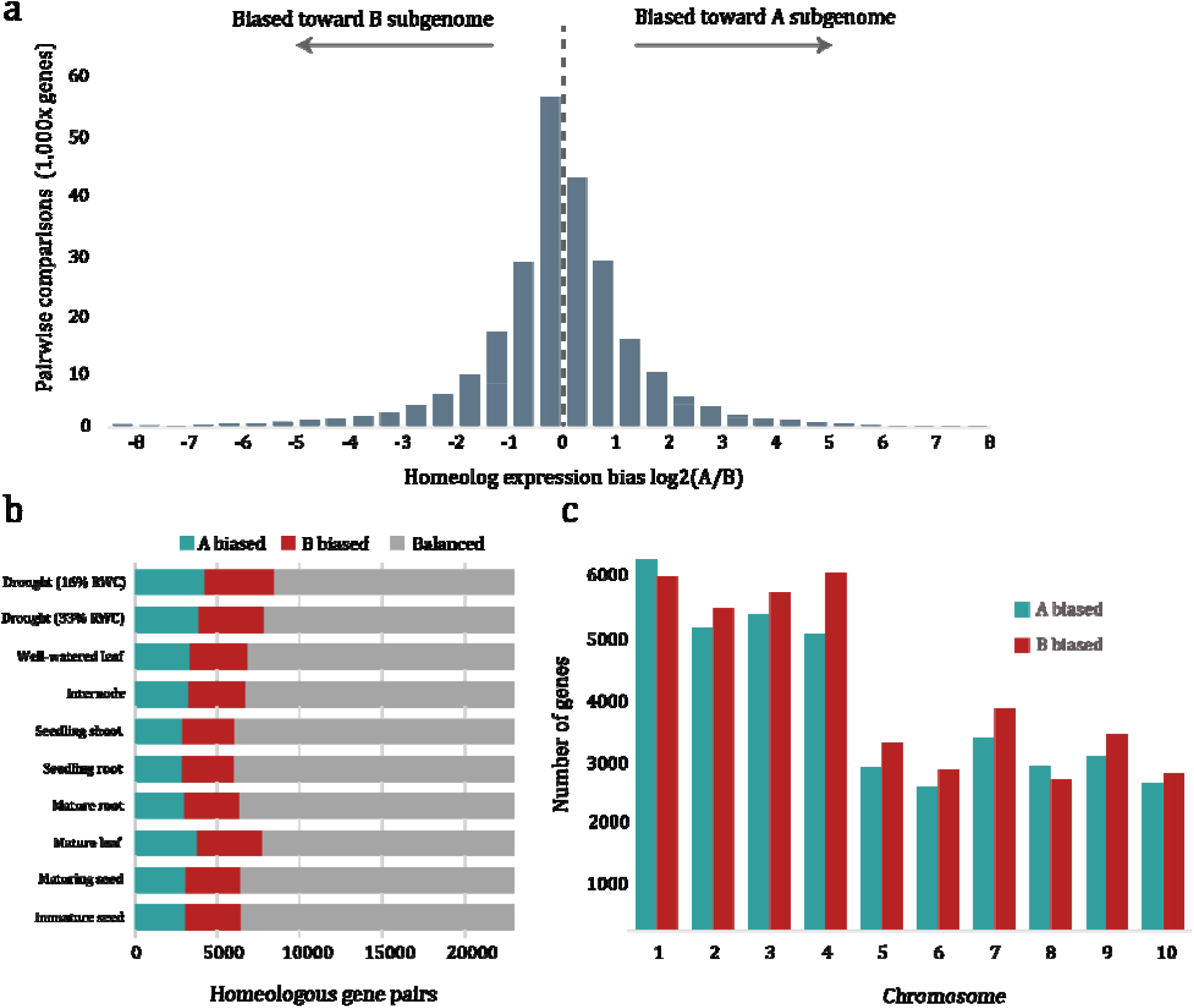
Homoeolog expression bias between the A and B subgenomes of teff. (a) The distribution of homoeolog expression bias (HEB) between all gene pairs in all tissues. An HEB > 0 indicates bias toward the A subgenome and a HEB < 0 indicates bias toward the B subgenome. (b) HEB across the ten tissues in the teff expression atlas. Gene pairs were classified as biased toward the A (blue) or B (red) subgenomes or balanced with no statistically significant differential expression (grey). (c) HEB in each of the ten pairs of chromosomes across all ten tissue types.

Teff belongs to the Chloridoideae subfamily of grasses ^27^ which includes important drought and heat tolerant C4 species such as the orphan grain crop finger millet and model desiccation tolerant plants in the genera *Oropetium, Eragrostis, Tripogon, Sporobolus*, and others. Most (∼90%) of surveyed Chloridoideae species are polyploid, including many of the aforementioned taxa, and this likely contributes to their diversity and stress tolerance ^16^. We utilized the wealth of genomic resources within Chloridoideae and more generally across Poaceae to identify patterns associated with improved stress tolerance, polyploidy and genome evolution in teff. The teff and Oropetium genomes have near complete collinearity, as demonstrated by highly conserved gene content and order along each chromosome (Figure 3). Teff and Oropetium show a clear 1:2 synteny pattern with 87% of teff genes having synteny to one block in Oropetium and 85% of Oropetium genes having synteny to two blocks in the teff genome (Figure 3a). This ratio corresponds to the A and B homoeologs of tetraploid teff and the single orthologs of diploid Oropetium. Each Oropetium chromosome has clear collinearity to two homoeologous teff chromosomes (Figure 3c). Three trios have no rearrangements (teff 3A, 3B, and Oropetium Chr3; 4A, 4B, Chr4; 6A, 6B, Chr8) six trios have one or more large-scale inversions (1A, 1B, Chr1; 2A, 2B, Chr2; 5A, 5B, Chr7; 7A, 7B, Chr6; 8A, 8B, Chr9; 9A, 9B, Chr5) and one trio has translocations (10A, 10B, Chr10). Of the 28,909 Oropetium genes, 74% (21,293) have syntenic orthologs in both subgenomes of teff, 5% (1,503) are found in only one subgenome, and 21% (6,113) have no syntenic orthologs in teff. Teff and the allotetraploid grain crop finger millet have 2:2 synteny but only 69% of syntenic blocks are found in duplicate because of the fragmented nature of the finger millet genome assembly ^28^ (Supplemental Figure 5). Only 56% (38,149) of the teff genes have two syntenic orthologs in finger millet and the remaining 13 and 30% (9,228 and 20,878) have one or zero syntenic orthologs in finger millet respectively.

Using Oropetium and teff syntenic orthologs, we calculated the ratio of nonsynonymous (Ka) to synonymous substitutions (Ks) to identify genes putatively under selection during domestication in teff. The top 10% of genes with the highest Ka/Ks ratios in teff (cutoff of 0.38) are enriched in gene ontology (GO) terms related to somatic embryogenesis, pollen differentiation, and reproductive phase transition among others (Supplemental Table 6). These genes may have been intentional or inadvertent targets during domestication.

Following an allopolyploidy event, a dominant subgenome often emerges with significantly more retained genes and higher homoeolog expression as the plant returns to a diploid-like state ^13^. This dominance is established immediately following the polyploidy event ^15^, and patterns of biased fractionation have been observed in Arabidopsis ^13^, maize ^29^, *Brassica rapa* ^30^, and bread wheat ^31^. Biased homoeolog loss (fractionation) is not universal, and other allopolyploids such as *Capsella bursa-pastoris* ^32^ and several Cucurbita species ^33^ display no subgenome dominance. We searched for biased fractionation using syntenic orthologs from Oropetium as anchors. The A and B subgenomes of teff have a near identical number of syntenic orthologs to Oropetium (19,277 vs. 19,292 respectively) suggesting that there is little or no biased fractionation in teff. Orthologs to 1,308 Oropetium genes are found as single copy loci in teff, including 647 and 678 from the A and B subgenomes respectively. The remaining orthologs are maintained in duplicate in teff compared to their single ortholog in Oropetium. Together this suggests a general stability of gene content in *Eragrostis* after genome merger.

### Homoeolog expression patterns and subgenome dominance

To test for patterns of sub-genome differentiation and dominance in teff, we surveyed gene expression in eight developmentally distinct tissue types and two stages of progressive drought stress. Sampled tissues include roots and shoots from seedlings and mature plants, internodes, and two stages of developing seeds. Tissue from mature, well-watered leaves and two time points of severe drought were also collected (leaf relative water content of 33% and 16% respectively). Of the 23,303 syntenic gene pairs between the A and B subgenome, 15,325 have homoeologous expression bias (HEB) in at least one tissue and 1,694 have biased expression in all sampled tissues (Supplemental Figure 6). Pairwise comparisons between syntenic gene pairs support a slight bias in transcript expression toward the B subgenome (Figure 4a). Roughly 56% of the 207,873 pairwise comparisons across the ten tissues show biased expression toward homoeologs in the B subgenome (Wilcoxon rank sum *P* < 0.001). This pattern is consistently observed across all ten tissues and most chromosome pairs, but the difference is subtle when robust cutoffs of differential expression are applied (Figure 4b and c; see methods). Individual tissues have from 6,061 to 8,485 homoeologous gene pairs with significant differential expression, including 52.3% biased toward the B subgenome (Kruskal–Wallis H test *P* < 0.01; Figure 4b). Eight pairs of chromosomes show HEB toward the B subgenome, and chromosomes 1 and 8 have more dominant homoeologs from the A subgenome, but the difference is not significant (Wilcoxon rank sum *P* > 0.05). Together this suggests that the B subgenome is universally dominant over the A subgenome but when strict thresholds are applied, this difference is minimal. Although we detected no evidence of recent homoeologous exchange, it is possible that genes from the recessive genome were replaced with homoeologs from the dominant subgenome, which would weaken patterns of subgenome dominance ^34^.

We tested whether gene pairs with HEB maintain patterns of dominance across all tissues or whether dominant homoeologs are reversed in different tissues or under stress. The vast majority of genes (86.9%; 13,322) with homoeologous expression bias maintain the same pattern of dominance across all tissues, while 13.1% (2,002) of gene pairs have opposite dominance patterns in different tissues. The remaining 7,675 gene pairs have no expression bias in any tissues or both homoeologs have negligible expression. Severely dehydrated leaf tissue had the most gene pairs with HEB (36%; 8,485) compared to seedling roots and shoots which each had ∼26% of pairs with HEB. These results are consistent with previous findings in allohexaploid wheat ^35^ and allotetraploid *Tragopogon mirus* ^36^. We compared the ratio of nonsynonymous (Ka) to synonymous substitution rates (Ks) in homoeologous gene pairs to test if genes with stronger HEB are experiencing different patterns of selection. Gene pairs with stronger HEB had significantly higher Ka/Ks than gene pairs with no HEB in any tissue (Supplemental Figure 7; 0.17 vs. 0.28; Mann-Whitney P < 0.01). We detected no difference in divergence (Ks) among genes with varying degrees of HEB (Supplemental Figure 8). This suggests homoeologous gene pairs with higher expression divergence are under more relaxed selective constraints than gene pairs with balanced expression.

## Discussion

Unlike the genomes of most polyploid grasses, the teff subgenomes are relatively small (∼300 Mb), with high gene density and low transposable element content. The subgenomes are highly syntenic along their length with no evidence of major inversions or structural rearrangements, contrasting patterns observed in other similarly aged allopolyploids such as wheat ^37^, canola (*Brassica napus*) ^38^, strawberry (*Fragaria ananassa*) ^39^, cotton ^40^, and proso millet ^41^. The general stability of the teff subgenomes may be attributed to low rates of homoeologous exchange. An estimated 90% of Chloridoid grasses are polyploid and among the allopolyploid species, multivalent pairing is rarely detected ^16^. The twenty chromosome pairs in teff show bivalent pairing in meiosis I ^22^, and double reduction has not been observed in segregating populations ^42,43^. Although homoeologous exchanges can result in advantageous emergent phenotypes, they can also destabilize the karyotype, leading to reduced fertility and fitness^44^. For this reason, recent polyploids have long been considered “evolutionary dead ends” ^45^. Thus, proper bivalent pairing (disomic inheritance) in natural allopolyploids may be favored, and the near perfect synteny observed between teff subgenomes suggests that an underlying mechanism may exist to prevent or reduce homoeologous exchanges in this species. We detected no evidence of recent homoeologous exchange in teff based on Ks distribution, including exchanges that would have happened at the inception of the polyploidy event 1.1 mya. Homeologous exchanges are a common feature of allopolyploids ^34^, and the lack of these events is a unique feature of the teff genome.

The Teff A and B subgenomes, and *Oropetium genome* have high degrees of chromosome level collinearity despite their distant divergence. This is particularly unusual as polyploidy-rich lineages typically have high rates of chromosome evolution ^46^. In contrast, our analysis of the divergence dates of the diploid A and B genome ancestors (∼5 mya) and the formation of the tetraploid (∼1.1 mya) indicates that the two genomes were so similar in structure (i.e., gene content, gene order and chromosome size) that some tetrasomic pairing would have been expected. Perhaps the status of the *Ph1*-equivalent locus (loci) ^47^ in *Eragrostis* is (are) so dominant, that even low frequencies of homoeologous pairing are blocked. The high levels of subgenome compatibility, genetic and chromosome stability, fidelity for chromosome pairing, and low rates of homoeologous exchange allows polyploidy to dominate in the Chloridioideae subfamily. This polyploidy in turn may have enabled the emergent resilience and robustness observed in Chloridoid grasses.

Although we detected no biased fractionation between the teff subgenomes, we observed a general subgenome dominance across tissues in the expression atlas. The B subgenome is smaller and has fewer transposable elements, which may be contributing to the overall higher homoeolog expression levels ^15^. Patterns of B subgenome dominance are relatively weak compared to other allopolyploids ^15^, which may reflect the stability and lack of biased fractionation in teff. The teff subgenomes have successfully partitioned their ancestral roles, and most gene pairs display homoeolog expression bias. This bias is generally maintained across tissues and treatments, and few gene pairs change bias in a tissue-specific manner. Severely drought stressed leaf tissue has the highest proportion of genes with biased expression, which may reflect adaptation to adverse environments. Extensive homoeolog expression bias is also observed in hexaploid wheat ^35^, octoploid sugarcane ^48^, and tetraploid *Tragopogon mirus* ^36^ and may be a common feature of recent polyploid grasses.

The vast majority of genes in Teff are maintained as homoeologous gene pairs in the A and B subgenomes, providing a significant obstacle for targeted breeding. Efforts to produce semi-dwarf, lodging resistant teff using a mutagenesis approach have been more difficult because of gene redundancy ^49^. The resources provided here will help accelerate marker-assisted selection and guide genome engineering-based approaches, which must take gene redundancy into account. Most gene pairs have divergent expression profiles such that the subgenomes likely contribute unequally to different agronomic traits. Teff is often described as an orphan grain crop because of its limited investigation and improvement, resulting in relatively low yields under ideal conditions compared to other cereals with intensive selection and breeding histories. Teff and other grasses within Chloridoideae have high tolerance to abiotic stresses, and most of this resilience was maintained during teff domestication. This may represent a historical alternative selection scheme where maximum yield is exchanged for reliable harvest under poor environmental conditions. Future efforts to improve food security should utilize the natural resilience of these robust, stable, polyploid species.

## Methods

### Plant materials

The ‘Dabbi’ cultivar of teff (PI 524434, www.ars-grin.gov) was chosen for sequencing and for constructing the expression atlas. Plant materials for High molecular weight (HMW) genomic DNA extraction, Hi-C library construction and RNA were maintained in growth chambers under a 12-hour photoperiod with day/night temperatures of 28°C and 22°C respectively and a light intensity of 400 μE m−2 sec−1. Tissue samples for the expression atlas were collected at ZT8 (Zeitgeber Time 8) to reduce issues associated with circadian oscillation. The tissue types used in the expression atlas include shoots and roots from young seedlings, mature leaf, internode, root, immature seeds and mature seeds. For the drought time points, mature teff plants were allowed to dry slowly and leaf tissue was collected at subsequent days of extreme drought when the plant tissues had 33% and 16% relative water content, as well as well-watered teff for comparison. Three biological replicates were collected for each sample in the expression atlas. Leaf tissue from seedlings was used for the HMW genomic DNA extraction and Hi-C library construction. Tissues for HMW genomic DNA extraction and RNAseq were immediately frozen in liquid nitrogen and stored at −80° C.

### DNA isolation, library construction, and sequencing

HMW genomic DNA was isolated from young teff leaf tissue for both PacBio and Illumina sequencing. A modified nuclei preparation ^50^ was used to extract HMW gDNA and residual contaminants were removed using phenol chloroform purification. PacBio libraries were constructed using the manufacturer’s protocol and were size selected for 30 kb fragments on the BluePippen system (Sage Science) followed by subsequent purification using AMPure XP beads (Beckman Coulter). The PacBio libraries were sequenced on a PacBio RSII system with P6C4 chemistry. In total, 5.5 million filtered PacBio reads were generated, collectively spanning 52.9 Gb or ∼85x genome coverage (assuming a genome size of 622 Mb). The same batch of HMW genomic DNA was used to construct Illumina DNAseq libraries for correcting residual errors in the PacBio assembly. Libraries were constructed using the KAPA HyperPrep Kit (Kapa Biosystems) followed by sequencing on an Illumina HiSeq4000 under paired-end mode (150 bp).

### RNA extraction and library construction

RNA for the expression atlas was extracted using the Omega Biotek E.Z.N.A. ® Plant RNA kit according to the manufacturer’s protocol. Roughly 200 mg of ground tissue was used for each extraction. The RNA quality was validated using gel electrophoresis and the Qubit RNA IQ Assay (ThermoFisher). Stranded RNAseq libraries were constructed using 2ug of total RNA quantified using the Qubit RNA HS assay kit (Invitrogen, USA) with the Illumina TruSeq stranded total RNA LT sample prep kit (RS-122-2401 and RS-122-2402). Multiplexed libraries were pooled and sequenced on an Illumina HiSeq4000 under paired-end 150nt mode. Three replicates were sequenced for each timepoint/sample.

### Genome assembly

The genome size of ‘Dabbi’ teff was estimated using flow cytometry as previously described ^51^. The estimated flow cytometry size was 622 Mb, which was consistent with kmer-based estimations from Illumina data. The kmer plot had a unimodal distribution suggesting low within genome heterozygosity and high differentiation from the teff A and B subgenomes. Raw PacBio data was error corrected and assembled using Canu (V1.4) ^52^ which produced accurate and contiguous assembly for homozygous plant genomes. The following parameters were modified: minReadLength=2000, GenomeSize=622Mb, minOverlapLength=1000. Assembly graphs were visualized after each iteration of Canu in Bandage ^53^ to assess complexities related to repetitive elements and homoeologous regions. The final Canu based PacBio assembly has a contig N50 of 1.55 Mb across 1,344 contigs with a total assembly size of 576 Mb. The raw PacBio contigs were polished to remove residual errors with Pilon (V1.22) ^18^ using 73x coverage of Illumina paired-end 150 bp data. Illumina reads were quality-trimmed using Trimmomatic ^54^ followed by aligning to the assembly with bowtie2 (V2.3.0) ^55^ under default parameters. Parameters for Pilon were modified as follows: --flank 7, --K 49, and --mindepth 15. Pilon was run recursively three times using the modified corrected assembly after each round. Ten full-length fosmids (collectively spanning 351kb) were aligned to the final PacBio assembly to assess the quality. The fosmids exhibited an average identity of 99.9% to the PacBio assembly, with individual fosmids ranging from 99.3 to 100% nucleotide identity.

### Hi-C analysis and pseudomolecule construction

The PacBio based teff contigs were anchored into a chromosome-scale assembly using a Hi-C proximity-based assembly approach as previously described ^19^. A Hi-C library was constructed using 0.2 g of leaf tissue collected from newly emerged teff seedlings with the Proximo™ Hi-C Plant kit (Phase Genomics) following the manufacturer’s protocol. After verifying quality, the Hi-C library was size-selected for 300-600 bp fragments and sequenced on the Illumina HiSeq 4000 under paired-end 150 bp mode. 150 bp reads were used to avoid erroneous alignment in highly similar homoeologous regions. In total, 226 million read pairs were used as input for the Juicer and 3d-DNA Hi-C analysis and scaffolding pipelines ^21,56^. Illumina reads were quality-trimmed using Trimmomatic ^54^ and aligned to the contigs using BWA (V0.7.16) ^20^ with strict parameters (-n 0) to prevent mismatches and non-specific alignments in repetitive and homoeologous regions. Contigs were ordered and oriented and assembly errors were identified using the 3d-DNA pipeline with default parameters ^56^. The resulting hic contact matrix was visualized using Juicebox, and misassemblies and misjoins were manually corrected based on neighboring interactions. This approach identified 20 high-confidence clusters representing the haploid chromosome number in Teff. The manually validated assembly was used to build pseudomolecules using the finalize-output.sh script from 3d-DNA and chromosomes were renamed and ordered by size and binned to the A and B subgenomes based on centromeric array analysis (described in detail below).

### Identification of repetitive elements

We first identified and masked the simple sequence repeats in the teff genome with GMATA ^57^, and then conducted structure-based full-length transposable element (TE) identification using the following bioinformatic tools: LTR_FINDER ^58^ and LTRharvest ^59^ to find LTR-RTs, LTR_retriever ^60^ to acquire high-confidence full LTR retrotransposons, SINE-Finder ^61^ to identify SINEs, MGEscan-nonLTR (V2) ^62^ to identify LINEs, MITE-Hunter ^63^ and MITE Tracker ^64^ to identify TIRs, and HelitronScanner ^65^ to identify *Helitrons*. All TEs were classified and manually checked according to the nomenclature system of transposons as described previously ^66^ and against Repbase to validate their annotation ^67^. We used the newly identified TEs as a custom library to identify full length and truncated TE elements through a homology-based search with RepeatMasker (http://www.repeatmasker.org, version 4.0.7) ^68^ using the teff pseudomolecules as input. Parameters for RepeatMasker were as described previously ^69^, and all other parameters were left as default. The distribution of repeat sequences was then calculated. Only LTR-RT families with at least 5 intact copies were used for analysis of subgenome specificity. Within the 65 families having > 5 intact elements, we identified LTRs with subgenomic specific activity. A family is considered as subgenomic specific if all intact elements of this family are from the same subgenome. Subgenome specificity was verified through BLAST of the element against the genome, and the distribution of matched sequences was manually inspected for subgenome specificity. The approximate insertion dates of LTR-RTs were calculated using the evolutionary distance between two LTR-RTs ^70,71^ with the formula of T=K/2μ, where K is the divergence rate approximated by percent identity and μ is the neutral mutation rate estimated as μ=1.3 × 10-8 mutations per bp per year.

Centromeric repeat arrays were identified with the approach outlined in ^72^ using Tandem repeat finder (Version 4.07) ^73^. Parameters were modified as follows for Tandem repeat finder: ‘1 1 2 80 5 200 2000 -d –h’. Centromere-specific repeats are often the most abundant tandem repeats in the genome, and they were identified in teff by the following criteria: (1) copy number, (2) sequence level conservation between chromosomes, (3) similarity to other grass repeats, and (4) proximity to centromere-specific *gypsy* LTR-RTs. This approach identified two distinct centromere-specific arrays (CenTA and CenTB) with a shared length of 159 bp yet distinct sequence compositions. The consensus sequence of centromeric repeats from each chromosome was used to construct a maximum likelihood phylogenetic tree implemented in MEGA5 (V10.0.5) ^74^. This approach separated centromeric repeats from the twenty chromosomes into two distinct groups corresponding to the A and B subgenomes.

### Genome annotation

Genes in the teff genome were annotated using the MAKER-P pipeline ^75^. The LTR-RT repeat library from LTR retriever was used for repeat masking. Transcript-based evidence was generated using RNAseq data from the ten tissues of the teff expression atlas. Quality trimmed RNAseq reads were aligned to the unmasked teff genome using the splice aware alignment program STAR (v2.6) ^76^ and transcripts were identified using StringTie (v1.3.4) ^77^ with default parameters. The –merge flag was used to combine the output from individual libraries to generate a representative set of non-redundant transcripts. Protein sequences from the Arabidopsis ^78^, rice ^79^, and sorghum ^80^ genomes as well as proteins from the UniProtKB plant databases ^81^ were used as protein evidence. *Ab initio* gene prediction was conducted using SNAP ^82^ and Augustus (3.0.2) ^83^ with two rounds of iterative training. The resulting gene models were filtered to remove any residual repetitive elements using BLAST with a non-redundant transposase library. The annotation quality was assessed using the benchmarking universal single-copy orthologs (BUSCO; v.2) ^84^ with the plant-specific dataset (embryophyta_odb9).

### RNAseq expression analysis and homoeolog expression bias

Gene expression levels were quantified with the pseudo-aligner Kallisto (v 0.44.0) ^85^ using the teff gene models as a reference. Paired-end Illumina reads from the ten tissues in the expression atlas were quality trimmed using Trimmomatic (V0.33) with default parameters and pseudo-aligned to the gene models with Kallisto under default parameters with 100 bootstraps per sample. The teff A and B subgenomes have high sequence divergence (∼7%) such that misalignment between homoeologs was minimal. Expression levels were quantified as Transcripts Per Million (TPM) and the three biological replicates were averaged for direct homoeolog comparisons.

### Comparative genomics

Homoeologous gene pairs between the teff A and B subgenomes and syntenic gene pairs across select grasses were identified using the MCSCAN toolkit implemented in python (https://github.com/tanghaibao/jcvi/wiki/MCscan-(Python-version)). Teff homoeologs were identified by all vs. all alignment using LAST, and hits were filtered using default parameters in MCSCAN with a minimum block size of 5 genes. This approach identified 23,303 homoeologous, syntenic gene pairs between the A and B subgenome. Homoeologs gene pairs with translocations were not identified using this syntenic approach and were thus excluded from analysis. Tandem gene duplicates in teff were identified from the all vs. all LAST output with a maximum gene distance of 10. Gene models from teff were aligned to the *Oropetium thomaeum* ^19,25^ and *Sorghum bicolor* ^80^ genes as outlined above for comparative genomics analyses across grasses. Macro and microsyntenic dot plots, block depths, and karyotype comparisons were generated in python using scripts from MSCAN.

Ka and Ks values were computed using a set of custom scripts available on GitHub: https://github.com/Aeyocca/ka_ks_pipe/. The homoeologous gene pair list from the teff subgenomes and syntenic orthologs between teff and Oropetium were used as input and the protein sequences from each gene pair were aligned using MUSCLE v3.8.31 ^86^. PAL2NAL (v14)^87^ was used to convert the peptide alignment to a nucleotide alignment and Ks values were computed between gene pairs using codeml from PAML (V4.9h) ^88^ with parameters specified in the control file found in the GitHub repository listed above.

### Data availability

The raw PacBio data, Illumina DNAseq, and RNAseq data are available from the National Center for Biotechnology Information Short Read Archive. RNAseq reads from the teff expression atlas were deposited to the National Center for Biotechnology Information Short Read Archive under bioproject PRJNA525065. The genome assembly and annotation for Tef is available from CoGe under genome ID: id50954.

## Acknowledgments

We are indebted to Tsegaye Dabi at the Salk Institute for Biological Studies for introducing us to this amazing plant, and for inspiring generations of plant biologists. We thank Elliott Meer for assistance with PacBio sequencing, and the Monsanto Genomics Team (Randy Kerstetter, Mitch Sudkamp, Phil Latreille, Zijin Du and Joe Zhou) for full length sequenced fosmids. We thank James Schnable for his helpful comments and suggestions on the manuscript. This work is supported by funding from the National Science Foundation (MCB-1817347 to R.V.), Department of Energy (DE-SC0012639 to T.C.M. and T.P.M.), and partial support from the Bill & Melinda Gates Foundation (T.C.M. and D.B.).

