## Supplemental Information for "Exceptional subgenome stability and functional divergence in allotetraploid teff, the primary cereal crop in Ethiopia"

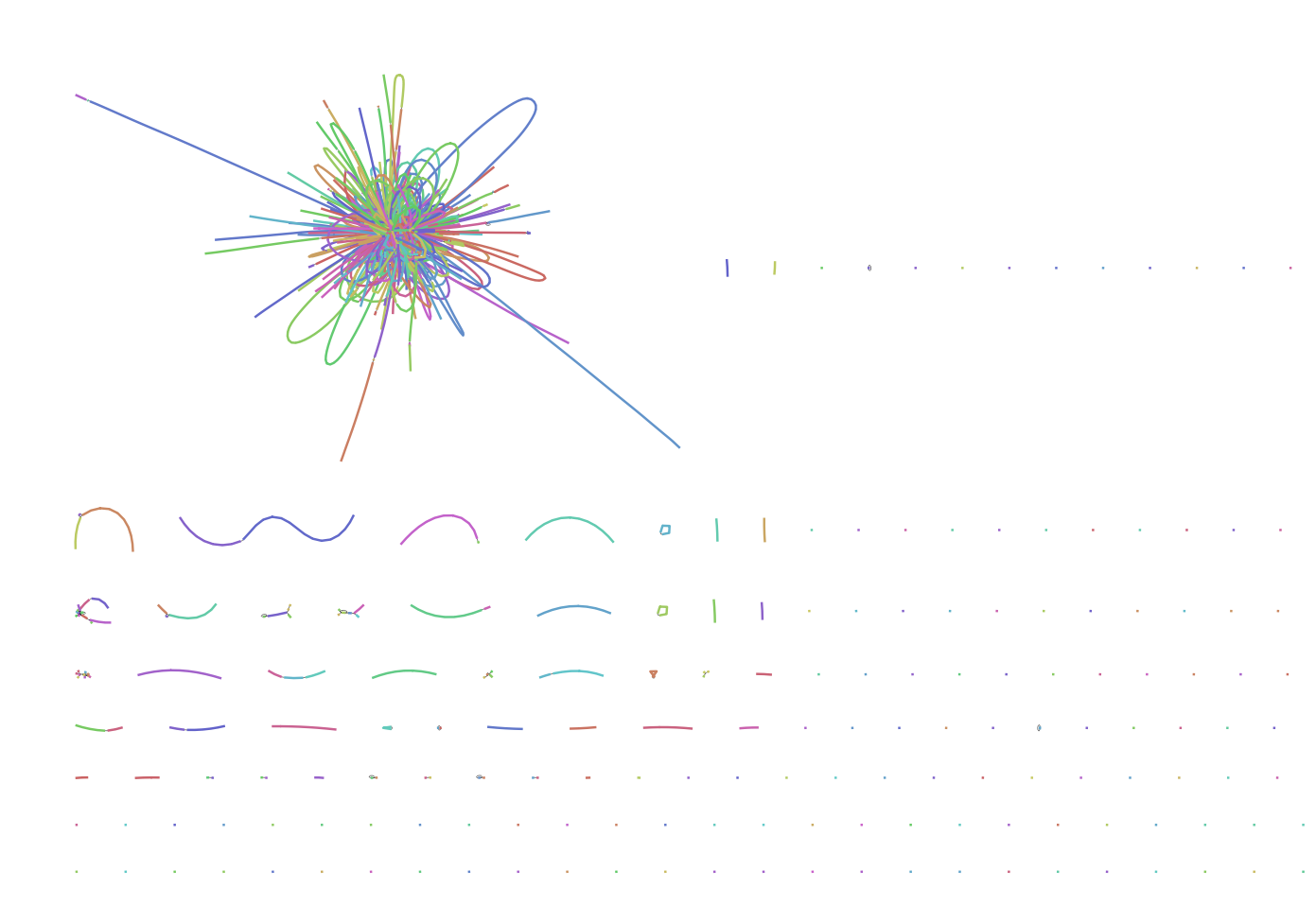

**Supplemental Figure 1. Genome assembly graph of the tef genome.** Each line represents a contig in the final Canu based assembly with connections representing ambiguities in the graph structure. The colors are assigned randomly.

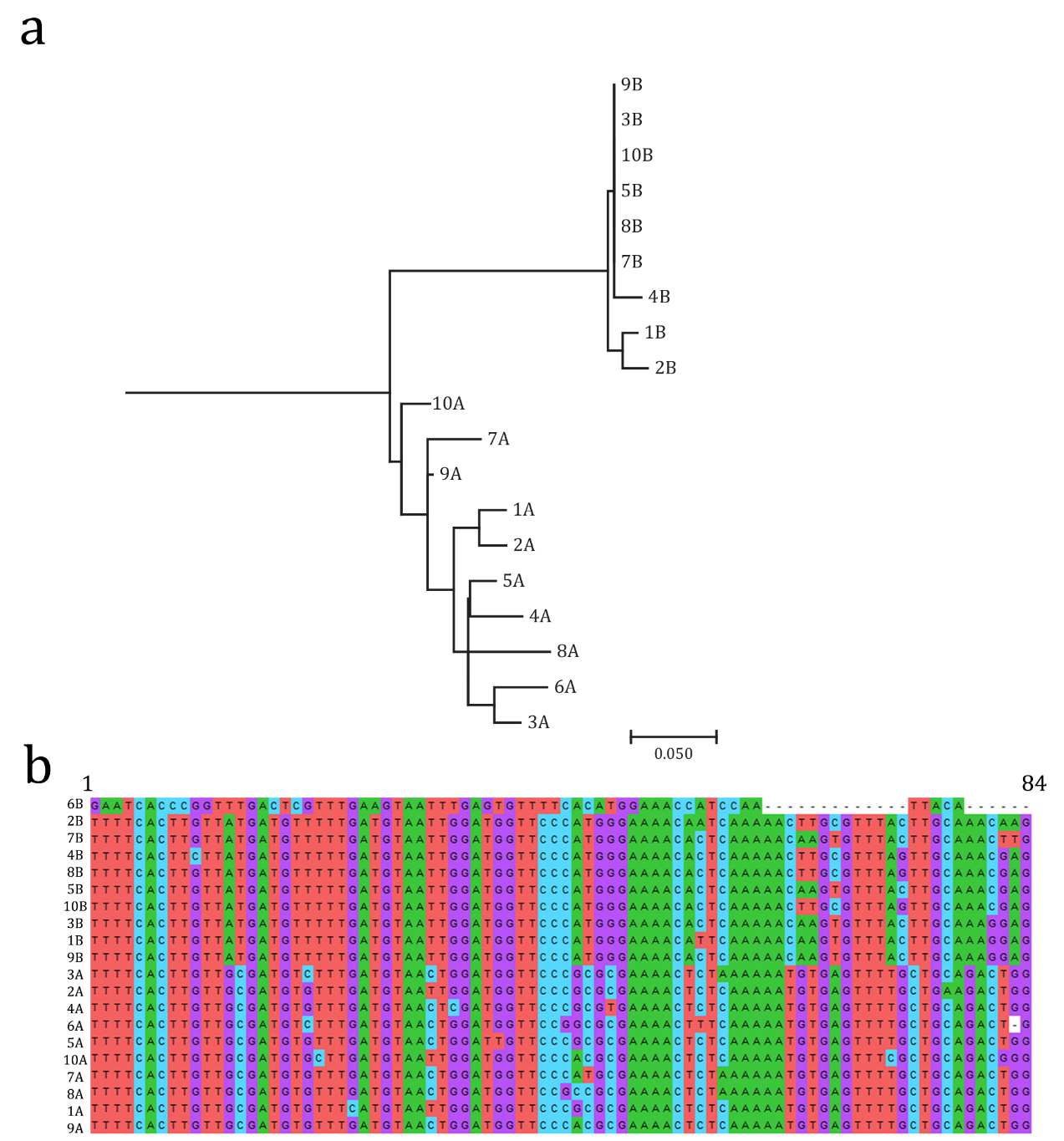

**Supplemental Figure 2. Classification of centromeric repeat arrays in the A and B subgenomes.** (a) Maximum likelihood phylogenetic tree of the consensus Cen sequence for each of the 20 chromosomes. (b) Alignment of the CenTA and CenTB repeat arrays. A 84 bp subset of the 159 bp Cen array with high homology is shown.

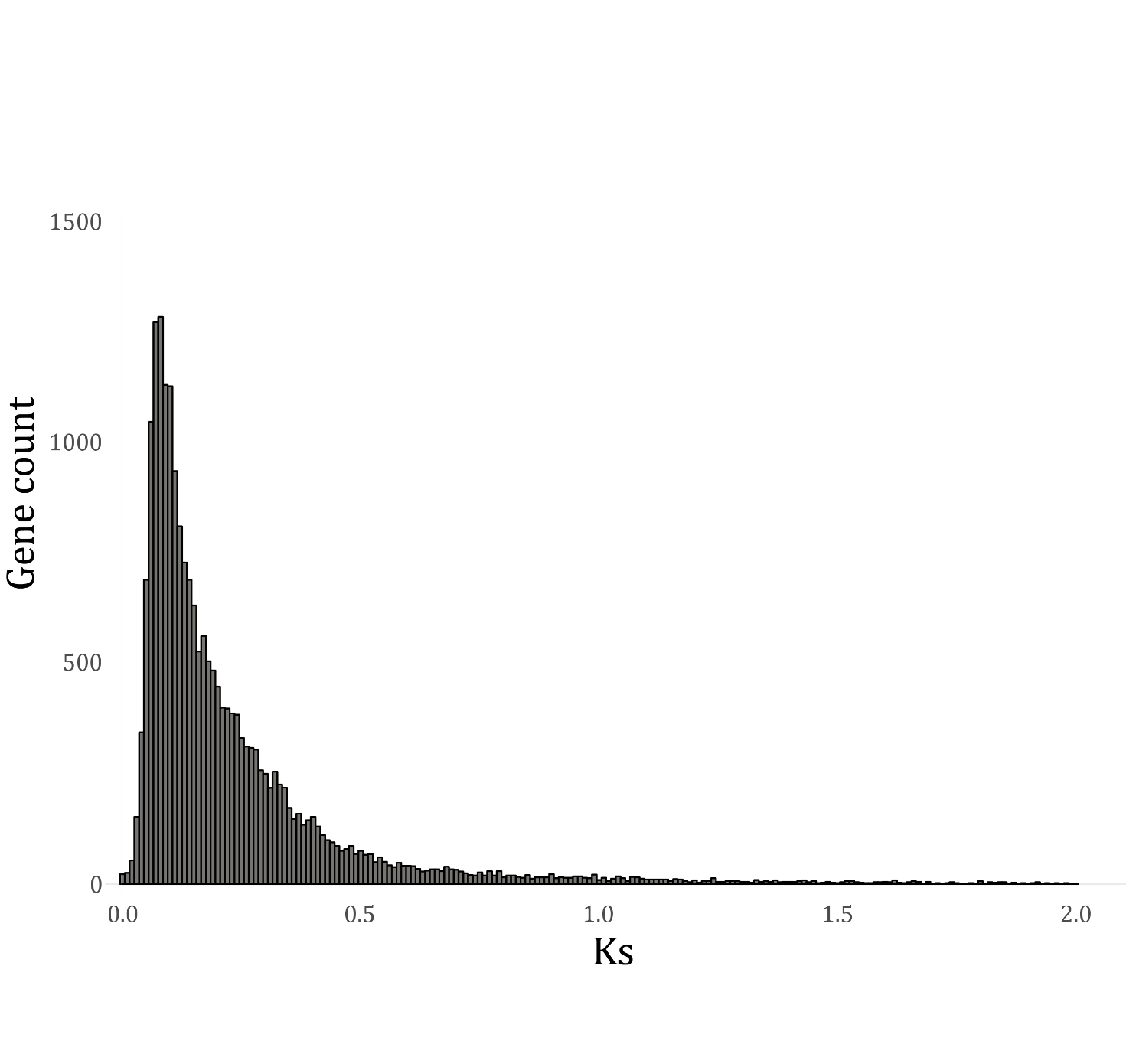

**Supplemental Figure 3.** Dating the divergence of the two subgenomes in tef. The distribution of Ks between homeologous gene pairs in the A and B subgenome is plotted.

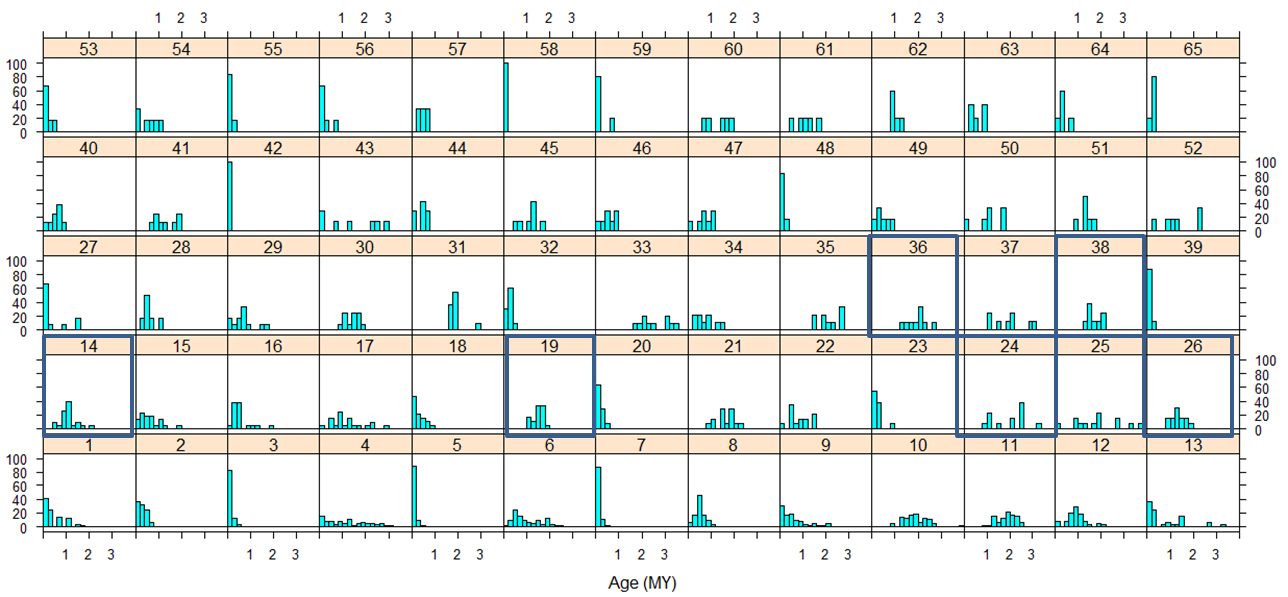

**Supplemental Figure 4. Histogram of insertion times of 64 LTR families that having >= 5 intact LTR elements.** In each panel, the Y-axis shows percentage and X-axis shows insertion time. The 6 subgenomic specific families are marked by blue blocks. Bin width = 0.2 MY.

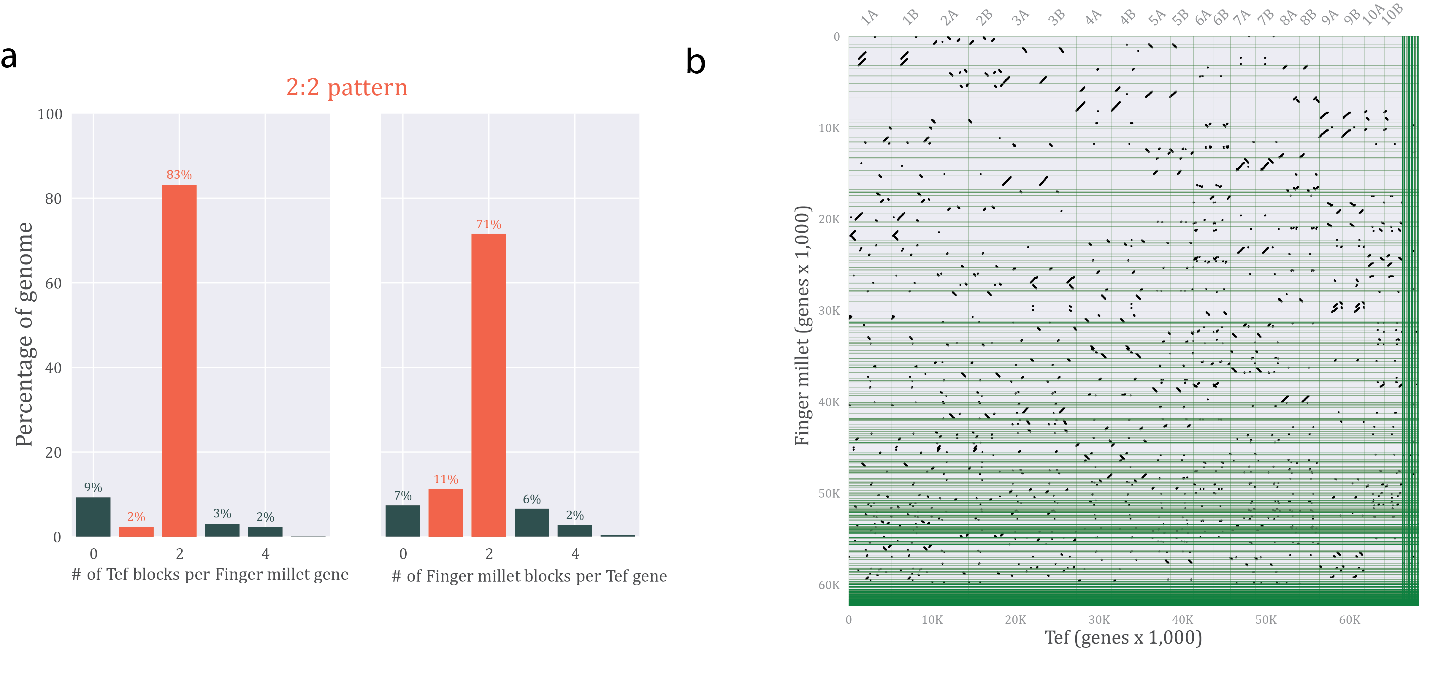

**Supplemental Figure 5. Comparative genomics between the allotetraploid tef and Finger millet genomes.** (a) Syntenic depth of tef blocks (left) and finger millet blocks (right) per finger millet and tef gene respectively. Syntenic blocks show a clear 2:2 pattern. (b) Macrosyntenic dotplot of the finger millet and tef genomes where each grey dot represents a syntenic gene pair.

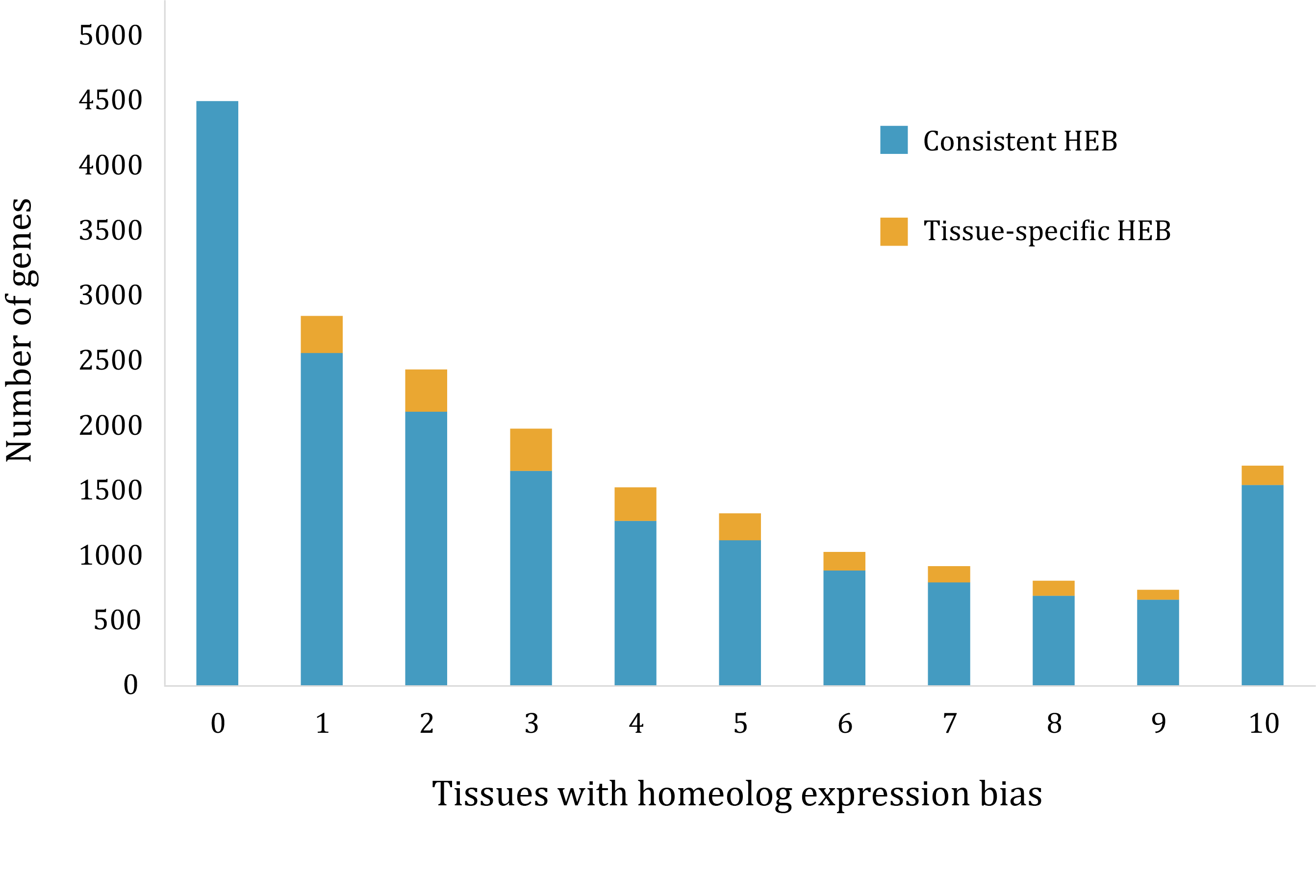

**Supplementary Figure 6. Conservation of homeolog expression bias across tissues.** A histogram showing the distribution of tissues showing homeolog expression bias (HEB) is plotted. Gene pairs showing consistent HEB are plotted in blue and genes with bias in both the A and B genomes in different tissues (tissue specific HEB) are shown in yellow.

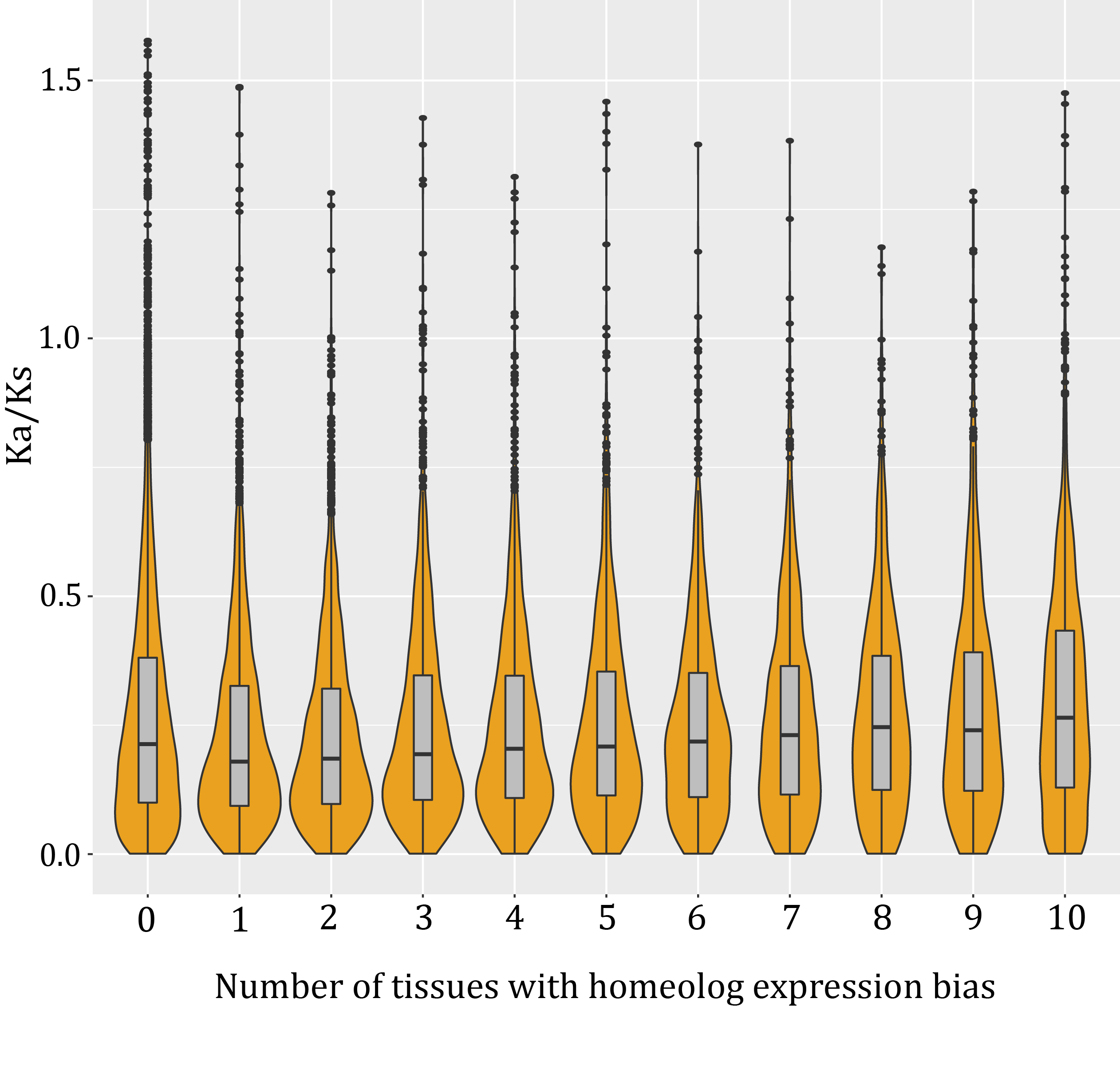

**Supplemental Figure 7. Patterns of selective constraint and homeolog expression bias (HEB).** Nested violin and box plots of Ka/Ks are shown for gene pairs ranging from 0 to 10 tissues with HEB.

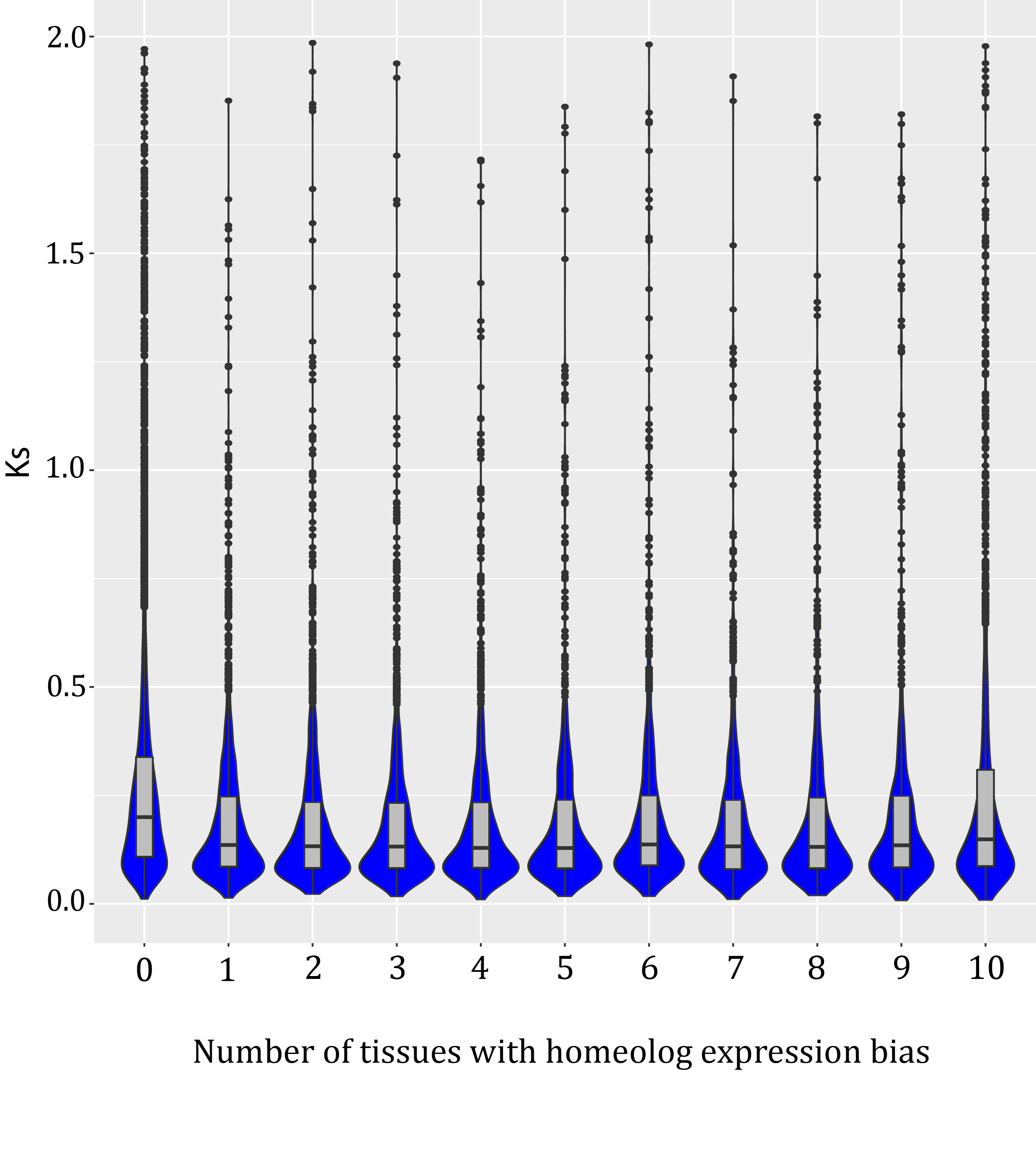

**Supplemental Figure 8. Divergence (Ks) of gene pairs and homeologous expression bias.** Nested violin and box plots of Ks are shown for gene pairs ranging from 0 to 10 tissues with HEB.

**Table 1.** Summary statistics of the tef genome

| **Chromosome** | **Size (bp)** | **Hi-C**  **Anchored contigs** | **Number of genes** | **Number of Tandem duplicates** |
| --- | --- | --- | --- | --- |
| 1A | 40,621,098 | 35 | 5,135 | 465 |
| 1B | 35,710,944 | 32 | 4,829 | 469 |
| 2A | 35,425,885 | 45 | 4,398 | 441 |
| 2B | 30,633,641 | 23 | 4,112 | 382 |
| 3A | 34,643,735 | 47 | 4,415 | 404 |
| 3B | 32,575,812 | 43 | 4,370 | 417 |
| 4A | 32,664,196 | 39 | 4,224 | 318 |
| 4B | 29,936,223 | 32 | 4,127 | 294 |
| 5A | 26,945,638 | 29 | 2,899 | 403 |
| 5B | 24,206,550 | 36 | 2,785 | 385 |
| 6A | 27,140,163 | 46 | 2,409 | 365 |
| 6B | 19,415,607 | 31 | 1,992 | 225 |
| 7A | 26,459,500 | 44 | 3,006 | 315 |
| 7B | 23,383,462 | 34 | 2,843 | 307 |
| 8A | 24,151,120 | 26 | 2,464 | 270 |
| 8B | 21,147,804 | 28 | 2,373 | 239 |
| 9A | 24,589,398 | 38 | 2,736 | 292 |
| 9B | 21,940,566 | 23 | 2,673 | 270 |
| 10A | 23,813,772 | 24 | 2,346 | 268 |
| 10B | 20,101,091 | 32 | 2,151 | 227 |
| unanchored | 22,232,506 | 657 | 1,968 | 130 |
| A subgenome | 296,454,505 | 373 | 34,032 | 3,541 |
| B subgenome | 259,051,700 | 314 | 32,255 | 3,215 |
| Total | 577,738,711 | 1,344 | 68,255 | 6,886 |

**Supplemental Table 1.** Summary of Fosmid alignment statistics to the tef genome

| **Fosmid** | **Tef Chromosome** | **Pos. start** | **Pos. end** | **Length (bp)** | **Mismatches (indels/SNPs)** | **% identity** |
| --- | --- | --- | --- | --- | --- | --- |
| 1 | Chromosome_2A | 24293453 | 24310772 | 17319 | 35 | 99.80 |
| 2 | Chromosome_3B | 480482 | 496500 | 16018 | 1 | 99.99 |
| 3 | Chromosome_1A | 607984 | 629933 | 21949 | 1 | 99.99 |
| 4 | Chromosome_2B | 1768602 | 1784819 | 16217 | 55 | 99.66 |
| 5 | Chromosome_6B | 9971798 | 9982526 | 10728 | 71 | 99.34 |
| 6 | Chromosome_7A | 18560345 | 18578197 | 17852 | 47 | 99.74 |
| 7 | Chromosome_1A | 607984 | 645075 | 37091 | 0 | 100 |
| 8 | Chromosome_2A | 13187152 | 13197778 | 10626 | 3 | 99.97 |
| 9 | Chromosome_1A | 629935 | 645075 | 15140 | 0 | 100 |
| 10 | Chromosome_8A | 7379205 | 7393614 | 14409 | 11 | 99.92 |
| 11 | Chromosome_3B | 467929 | 480309 | 12380 | 0 | 100 |
| 12 | Chromosome_6B | 19022180 | 19030707 | 8527 | 0 | 100 |
| 13 | Chromosome_3A | 15765021 | 15822822 | 57801 | 77 | 99.87 |
| 14 | Chromosome_8B | 2395467 | 2406855 | 11388 | 6 | 99.95 |
| 15 | Chromosome_2A | 25337979 | 25354980 | 17001 | 0 | 100 |
| 16 | Chromosome_4B | 4698711 | 4709012 | 10301 | 1 | 99.99 |
| 17 | Chromosome_7B | 1858973 | 1878530 | 19557 | 58 | 99.70 |
| 18 | Chromosome_3A | 23352162 | 23364268 | 12106 | 1 | 99.99 |
| 19 | Chromosome_5A | 18308369 | 18320474 | 12105 | 30 | 99.75 |
| 20 | Chromosome_1A | 35965810 | 35978550 | 12740 | 14 | 99.89 |
|  |  |  | **Total** | **351255** | **411** | **99.9** |

**Supplemental Table 2.** Summary ofcentromeric repeat array (CenT) composition in the tef genome

| **Chromosome** | **CenT Array start** | **CenT Array End** | **CenT Array Length (bp)** | **Number of CenT repeats** |
| --- | --- | --- | --- | --- |
| 1A | 21,557,537 | 21,623,717 | 66,180 | 399 |
| 1B | 19,501,629 | 19,520,995 | 19,366 | 114 |
| 2A | 14,451,025 | 14,548,454 | 97,429 | 569 |
| 2B | 12,104,147 | 12,108,587 | 4,440 | 26 |
| 3A | 14,292,984 | 14,296,855 | 3,871 | 24 |
| 3B | 14,092,648 | 14,176,019 | 83,371 | 121 |
| 4A | 17,363,887 | 17,401,206 | 37,319 | 99 |
| 4B | 15,985,907 | 16,152,093 | 166,186 | 824 |
| 5A | 15,924,190 | 15,941,051 | 16,861 | 103 |
| 5B | 13,811,928 | 13,872,280 | 60,352 | 115 |
| 6A | 12,945,587 | 13,004,019 | 58,432 | 348 |
| 6B | 8,377,013 | 8,380,775 | 3,762 | 22 |
| 7A | 21,934,795 | 22,036,202 | 101,407 | 449 |
| 7B | 21,827,654 | 21,831,745 | 4,091 | 24 |
| 8A | 14,254,377 | 14,301,695 | 47,318 | 398 |
| 8B | 12,502,475 | 12,627,067 | 124,592 | 117 |
| 9A | 9,554,342 | 9,614,054 | 59,712 | 189 |
| 9B | 7,617,457 | 7,629,471 | 12,014 | 71 |
| 10A | 13,918,716 | 14,165,131 | 246,415 | 347 |
| 10B | 10,984,530 | 11,310,738 | 326,208 | 231 |

**Supplemental Table 3.** Summary of the repeat sequence distribution

| **Class** | **Subclass** | **Superfamily** | **Family** | **Loci** | **Size (Mb)** | **Genome Size** |
| --- | --- | --- | --- | --- | --- | --- |
| SSR | SSR | unknown | 1 | 116936 | 5.2 | 0.9 |
| Class I | LTR | Gypsy | 944 | 54384 | 71.8 | 12.4 |
|  | LTR | unknown | 946 | 55889 | 32.5 | 5.6 |
|  | LTR | Copia | 330 | 13571 | 11.6 | 2 |
|  | LINE | L1 | 37 | 2784 | 1.6 | 0.3 |
|  | LINE | I | 5 | 17 | 0 | ~0 |
|  | SINE | Unknown | 109 | 14909 | 2.4 | 0.4 |
| Class II | TIR | Tc1 | 793 | 81715 | 14.9 | 2.6 |
|  | TIR | CACTA | 266 | 25197 | 4.4 | 0.8 |
|  | TIR | hAT | 77 | 7084 | 1.4 | 0.2 |
|  | TIR | PIF | 48 | 5746 | 1.1 | 0.2 |
|  | TIR | Mutator | 26 | 3238 | 0.6 | 0.1 |
|  | TIR | Unknown | 1 | 247 | 0 | ~0 |
|  | Helitron | Helitron | 105 | 21977 | 5.6 | 1 |
|  | Total |  |  |  | 153.1 | 26.5 |

**Supplemental Table 4.** Subgenome specificity of LTR-Retrotransposon

| **Famly ID** | **Subgenome** | **No. of intact**  **LTR-RT** | **Mean age** | **SD** | **Q25** | **Median age**  **(MYA)** | **Q75** |
| --- | --- | --- | --- | --- | --- | --- | --- |
| 14 | A | 23 | 1.1 | 0.3 | 1.0 | 1.1 | 1.2 |
| 19 | A | 18 | 1.5 | 0.2 | 1.3 | 1.5 | 1.7 |
| 24 | A | 13 | 2.0 | 0.7 | 1.2 | 2.2 | 2.4 |
| 26 | B | 13 | 1.3 | 0.3 | 1.1 | 1.4 | 1.5 |
| 36 | A | 9 | 2.0 | 0.4 | 1.7 | 2.1 | 2.1 |
| 38 | A | 8 | 1.7 | 0.3 | 1.5 | 1.6 | 2.0 |

**Supplemental Table 5.** Summary of LTR-Retrotransposon subgenome specificity

| **Category** | **No. of families** |
| --- | --- |
| Single burst < 1 MYA | 29 |
| Not showing burst, active period upper bound < 1 MY | 6 |
| Not showing burst, active period upper bound > 1 MY | 4 |
| Two bursts, one < 1MY, the other > 1 MY | 2 |
| Not showing burst, active period upper bound >= 2 MY | 15 |
| Single burst | 3 |

**Supplemental Table 6.** GO enrichment of the top 10% of genes with highest Ka/Ks ratios in tef compared to Oropetium

| Gene ontology | Description |
| --- | --- |
| GO:0006355 | regulation of transcription |
| GO:0006351 | transcription, DNA-templated |
| GO:0031627 | telomeric loop formation |
| GO:0010262 | somatic embryogenesis |
| GO:0090502 | RNA phosphodiester bond hydrolysis |
| GO:0035067 | negative regulation of histone acetylation |
| GO:0048235 | pollen sperm cell differentiation |
| GO:0033169 | histone H3-K9 demethylation |
| GO:0010228 | vegetative to reproductive phase transition |
| GO:0031054 | pre-miRNA processing |
| GO:0080188 | RNA-directed DNA methylation |
